# Identification of Drugs Blocking SARS-CoV-2 Infection using Human Pluripotent Stem Cell-derived Colonic Organoids

**DOI:** 10.1101/2020.05.02.073320

**Authors:** Xiaohua Duan, Yuling Han, Liuliu Yang, Benjamin E. Nilsson-Payant, Pengfei Wang, Tuo Zhang, Jenny Xiang, Dong Xu, Xing Wang, Skyler Uhl, Yaoxing Huang, Huanhuan Joyce Chen, Hui Wang, Benjamin tenOever, Robert E. Schwartz, David. D. Ho, Todd Evans, Fong Cheng Pan, Shuibing Chen

## Abstract

The current COVID-19 pandemic is caused by SARS-coronavirus 2 (SARS-CoV-2). There are currently no therapeutic options for mitigating this disease due to lack of a vaccine and limited knowledge of SARS-CoV-2 biology. As a result, there is an urgent need to create new disease models to study SARS-CoV-2 biology and to screen for therapeutics using human disease-relevant tissues. COVID-19 patients typically present with respiratory symptoms including cough, dyspnea, and respiratory distress, but nearly 25% of patients have gastrointestinal indications including anorexia, diarrhea, vomiting, and abdominal pain. Moreover, these symptoms are associated with worse COVID-19 outcomes^1^. Here, we report using human pluripotent stem cell-derived colonic organoids (hPSC-COs) to explore the permissiveness of colonic cell types to SARS-CoV-2 infection. Single cell RNA-seq and immunostaining showed that the putative viral entry receptor ACE2 is expressed in multiple hESC-derived colonic cell types, but highly enriched in enterocytes. Multiple cell types in the COs can be infected by a SARS-CoV-2 pseudo-entry virus, which was further validated *in vivo* using a humanized mouse model. We used hPSC-derived COs in a high throughput platform to screen 1280 FDA-approved drugs against viral infection. Mycophenolic acid and quinacrine dihydrochloride were found to block the infection of SARS-CoV-2 pseudo-entry virus in COs both *in vitro* and *in vivo*, and confirmed to block infection of SARS-CoV-2 virus. This study established both *in vitro* and *in vivo* organoid models to investigate infection of SARS-CoV-2 disease-relevant human colonic cell types and identified drugs that blocks SARS-CoV-2 infection, suitable for rapid clinical testing.

---

Previously, we reported a chemically-defined protocol to derive COs from hPSCs^2^, which we modified slightly based on published studies^3^. In brief, HUES8 hESCs were induced with CHIR99021 (CHIR) and Activin A to generate definitive endoderm (DE) (**Extended Data Fig. 1a**). After 4 days of culture with CHIR +FGF4 to induce hindgut endoderm (HE), cells were treated with BMP2, epidermal growth factor (EGF), and CHIR for 3 days to promote specification of colon progenitors (CPs). Starting on day 11, CPs were treated with a colonic medium containing CHIR, LDN193189 (LDN), and EGF. After embedding these organoids in Matrigel, spheroids became pseudostratified and progressively cavitated into fully convoluted organoids (**Fig. 1a**). The organoids expressed CDX2, Villin and SATB2, confirming colonic identity (**Fig. 1b**). Immunocytochemistry confirmed that COs contain cell types found in normal colon, including keratin 20 (KRT20)^+^ epithelial cells, mucin 2 (MUC2)^+^ goblet cells, EPH receptor B2 (EPHB2)^+^ transit-amplifying (TA) cells, and chromogranin A (CHGA)^+^ neuroendocrine (NE) cells (**Fig. 1c**).

**Figure 1.**
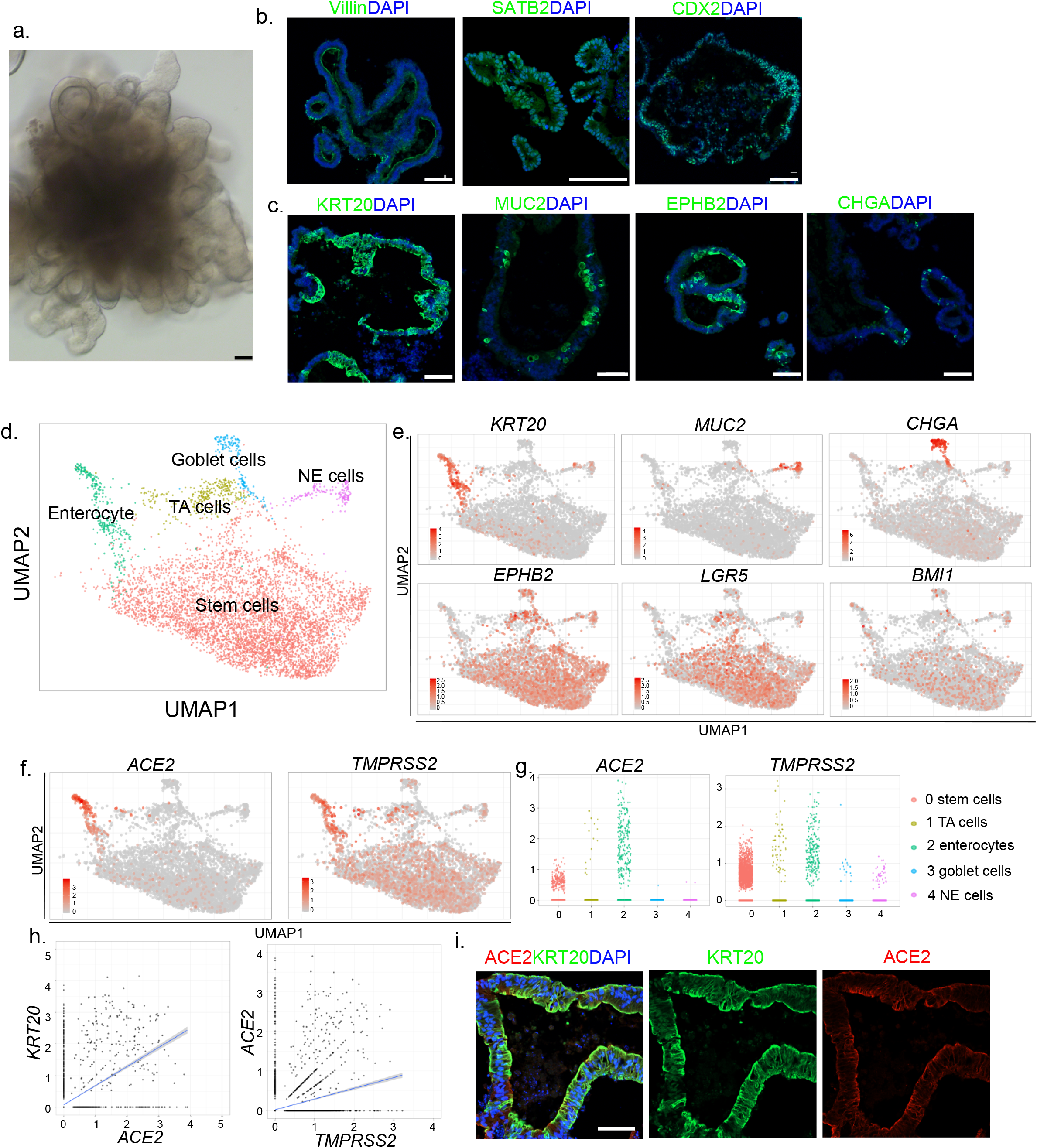
ACE2 and TMPRSS2 are expressed in hPSC-COs. **a,** Phase contrast image of a representative hPSC-derived colon organoid (CO). Scale bar = 100 μm. **b, c,** Confocal imaging of hPSC-COs stained with antibodies against (b) markers for colon cell fate, including Villin, SATB2, CDX2, or (c) KRT20, MUC2, EPHB2, and CHGA; DAPI stains nuclei. Scale bar = 100 μm. **d,** UMAP of hPSC-CO cell types. **e,** UMAP of markers for specific colonic cell fates markers, including *KRT20*, *MUC2*, *CHGA*, *EPHB2*, *LGR5*, and *BMI1*. **f,** UMAP of *ACE2* and *TMPRSS2*. **g,** Jitter plots for expression levels of *ACE2* and *TMPRSS2*. **h,**2D correlation of expression levels for *KRT20* and *ACE2*; *ACE2* and *TMPRSS2*. **i,** Representative confocal images of hPSC-COs co-stained with antibodies recognizing ACE2 and KRT20. DAPI stains nuclei. Scale bar = 100 μm.

Single cell RNA-seq was used to examine global transcript profiles at single cell resolution (**Extended Data Fig. 1b**). Consistent with the immunostaining results, most cells express *CDX2* and *VIL1* (**Extended Data Fig. 1c**). Five cell clusters were identified including *KRT20^+^* epithelial cells, *MUC2^+^* goblet cells, *EPHB2^+^* TA cells, *CHGA^+^* NE cells, and *LGR5^+^* or *BMI1^+^* stem cells (**Fig. 1d-e, Extended Data Fig. 1d**). We examined the expression of two factors associated with SARS-CoV-2 cell entry, the putative receptor *ACE2* and the protease *TMPRSS2*^4^. Both are expressed in all five cell clusters, but highly enriched in *KRT20^+^* enterocytes (**Fig. 1f-g**). Two-dimensional correlation confirmed the co-expression relationship for *ACE2* and *KRT20*, as well as *ACE2* and *TMPRSS2* (**Fig. 1h**). Immunohistochemistry further validated the co-expression of KRT20 and ACE2 in hPSC-COs (**Fig. 1i**).

To model infection of hPSC-COs with SARS-CoV-2, we used a vesicular stomatitis virus (VSV) based SARS-CoV-2 pseudo-entry virus, with the backbone provided by a VSV-G pseudo-typed ΔG-luciferase virus and the SARS-CoV-2 spike protein incorporated into the surface of the viral particle (See Methods for details)^5,6^. COs were fragmentized and innoculated with the SARS-CoV-2 pseudo-entry virus. 24 or 48 hr post-infection (hpi), the cells were lysed and monitored for luciferase activity (**Extended Data Fig. 2a**). The organoids infected with SARS-CoV-2 pseudo-entry virus at MOI=0.01 showed a strong signal at 24 hpi (**Fig. 2a**). Single cell RNA-seq was performed to examine the hPSC-derived COs at 24 hpi. The same five cell populations were identified in the COs post-infection (**Fig. 2b and Extended Data Fig. 2b-d**). Compared to uninfected samples, the *KRT20^+^* enterocyte population decreased significantly (**Fig. 2c**). Immunostaining confirmed increased cellular apoptosis, suggesting toxicity for these cells (**Extended Data Fig. 2e**). In addition, the *ACE2^+^* population was significantly depleted (**Fig. 2e**). The mRNAs of SARS-CoV-2 pseudo-entry virus, including *VSV-NS*, *VSV-N*, and *VSV-M*, were detected in all five cell populations (**Fig. 2f**), but not in the uninfected COs (**Extended Data Fig. 2f**). Immunostaining further validated the expression of luciferase in *ACE2^+^*, *VIL1^+^*, *CDX2^+^*, *KRT20^+^,* and *MUC2^+^* cells (**Fig. 2g**).

**Figure 2.**
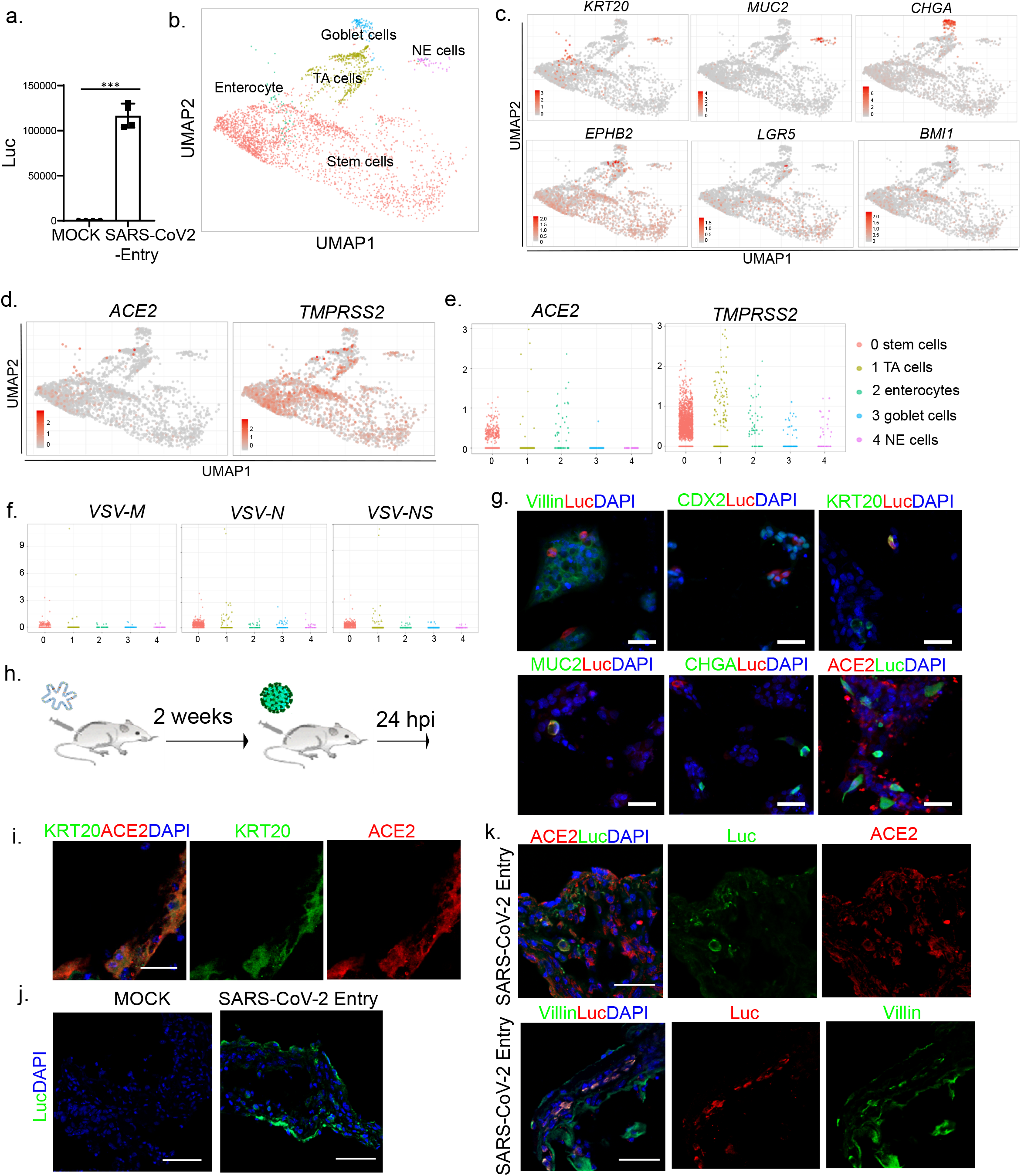
hPSC-COs can be infected by a SARS-CoV-2 pseudo-entry virus both *in vitro* and *in vivo*. **a,** Luciferase activity in lysates from hPSC-derived colonic cells at 24 hpi following exposure to SARS-CoV-2 pseudo-entry virus at MOI=0.01. **b,** UMAP of hPSC-COs at 24 hpi with SARS-CoV-2 pseudo-entry virus. **c,** UMAP of markers for specific colonic cell fates, including *KRT20*, *MUC2*, *CHGA*, *EPHB2*, *LGR5*, and *BMI1*. **d,** UMAP of *ACE2* and *TMPRSS2*. **e,** Jitter plots of *ACE2* and *TMPRSS2* transcript levels. **f,** Jitter plots of *VSV-M, VSV-N* and *VSV-NS* transcript levels. **g,** Immunocytochemistry staining of hPSC-CO cells infected with SARS-CoV-2 pseudo-entry virus (MOI=0.01) at 24 hpi using antibodies against luciferase and specific markers for colonic cell fate, including Villin, CDX2, KRT20, MUC2, CHGA, and ACE2. Scale bar= 50 μm. **h,** Schematic for the *in vivo* infection. **i,** Representative confocal image of a hPSC-CO xenograft at 24 hpi stained with antibodies against ACE2 and KRT20. DAPI stains nuclei. Scale bar = 25 μm. **j,** Representative confocal image of a hPSC-CO xenograft at 24 hpi (1×10^3^ FFU) stained with antibody against luciferase. DAPI stains nuclei. Scale bar = 75 μm. **k,** Representative confocal images of hPSC-CO xenografts at 24 hpi stained with antibodies against luciferase, ACE2 and Villin. DAPI stains nuclei. Scale bar = 50 μm. Data in (a) is presented as mean ± STDEV. *N=3. P* values were calculated by unpaired two-tailed Student’s t-test. \*\*\**P* < 0.001.

Humanized mice carrying hPSC-COs *in vivo* provide a unique platform for modeling COVID-19. In brief, hPSC-COs were transplanted under the kidney capsule of *NOD-scid IL2Rg^null^* mice. Two weeks after transplantation, the organoid xenograft was removed and examined for cellular identities (**Fig. 2h**). Consistent with *in vitro* culture, ACE2 can be detected in hPSC-derived KRT20^+^ enterocytes (**Fig. 2i**). SARS-CoV-2 pseudo-entry virus was inoculated locally. At 24 hpi, the xenografts were removed and analyzed by immuno-histochemistry. Luciferase was detected in the xenografts inoculated with virus, but not in MOCK-infected controls (**Fig. 2j**). Immunohistochemistry detected luciferase in ACE2^+^ and Villin^+^ cells, suggesting these are permissive to SARS-CoV-2 pseudo-entry virus infection *in vivo* (**Fig. 2k**).

Next, we adapted hPSC-COs to a high throughput screening platform and probed the Prestwick FDA-approved drug library to identify drug candidates capable of blocking SARS-CoV-2 pseudo-virus infection. In brief, hPSC-COs were cultured in 384-well plates. After overnight incubation, organoids were treated with drugs from the library at 10 μM. One hour post-exposure with drugs, the organoids were innoculated with the SARS-CoV-2 pseudo-entry virus. 24 hpi, the organoids were analyzed for luciferase activity (**Fig. 3a**). Drugs that decreased the luciferase activity by at least 75% were chosen as primary hit drugs (**Fig. 3b**). Eight drugs (**Extended Data Table 1**) were identified as lead hits and further tested for their capacities to decrease the luciferase signal in a dose-dependent manner (**Extended Data Fig. 3**). These drugs could potentially function through blocking virus entry, by decreased cell survival, or even by directly inhibiting luciferase activity. To distinguish these possibilities, the lead hit drugs were tested in comparison to hPSC-COs infected with a control VSVG-luciferase reporter virus. Four of the lead hit drugs showed specificity to SARS-CoV-2 pseudo-entry virus, including mycophenolic acid (MPA, **Fig. 3c**) (SARS-CoV-2: IC_50_=0.54 μM; VSVG: IC_50_=6.4 μM), quinacrine dihydrochloride (QNHC, **Fig. 3d**) (SARS-CoV-2: IC_50_=1.6 μM; VSVG: IC_50_=5.5 μM), chloroquine (SARS-CoV-2: IC_50_=6.8 μM; VSVG: IC_50_=19.1 μM), and resveratrol (SARS-CoV-2: IC_50_=5.7 μM; VSVG: IC_50_=39.3 μM) (**Fig. 3e-g,** and **Extended Data Fig. 3**). The IC_50_ of MPA is 10 times lower and the IC_50_ of QNHC is 5 times lower than that of chloroquine, a drug recently authorized by the FDA for emergency use to treat COVID-19 patients^7^. Immunostaining confirmed few luciferase positive cells in hPSC-COs treated with 3 μM MPA or 4.5 μM QNHC at 24 hpi (**Fig. 3h**). CO-explanted humanized mice were treated with 50 mg/kg MPA by IP injection, followed by local inoculation of SARS-CoV-2 pseudo-entry virus. At 24 hpi, the mice were euthanized, and xenografts were analyzed by immunostaining. Luc^+^ cells in the xenografts of MPA-treated mice were significantly lower than those of vehicle-treated mice (**Fig. 3i-k**).

**Figure 3.**
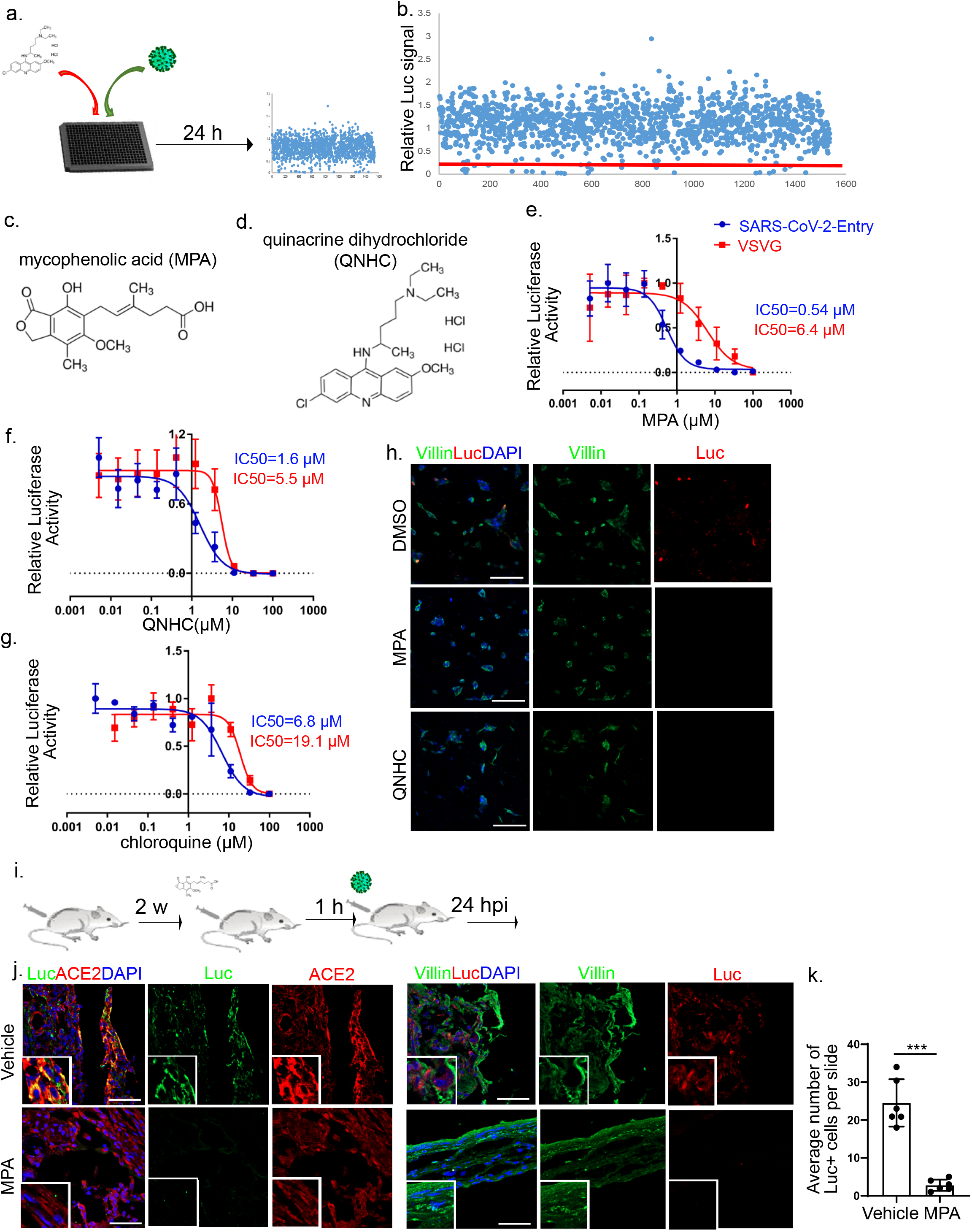
A high throughput screen to identify drugs that block SARS-CoV-2 entry. **a,** Schematic describing the high throughput drug screen platform. **b,** Primary screen data. **c, d,** Chemical structure of mycophenolic acid (MPA, c) and quinacrine dihydrochloride (QNHC, d). **e-g,** Inhibition curves for MPA (e), QNHC (f) and chloroquine (g) comparing hPSC-COs infected with SARS-CoV-2 pseudo-entry or control VSVG viruses (for both, MOI= 0.01). **h,** Immunofluorescent staining for luciferase in 3 μM MPA, 4.5 μM QNHC or DMSO-treated hPSC-COs at 24 hpi (MOI=0.01). Scale bar = 100 μm. **i.** Scheme of the *in vivo* drug evaluation. **j.** Representative confocal images of MPA or DMSO-treated hPSC-CO xenografts at 24 hpi stained with antibodies against luciferase, ACE2 and Villin. DAPI stains nuclei. Scale bar = 50 μm. **k.** Quantification of the average number of Luc^+^ cells per xenograft. (*N=6* xenografts, 5 slides/xenografts).

Finally, hPSC-COs were infected with SARS-CoV-2 virus at MOI=0.1 or 0.01. At 24 hpi, immunostaining detected the expression of SARS-CoV membrane protein in the infected hPSC-COs, which partially co-localized with CDX2 and KRT20 (**Fig. 4a**). Bulk RNA sequencing confirmed viral transcripts in the SARS-CoV-2 infected hPSC-COs (MOI=0.1, **Fig. 4b**). The MOCK and infected hPSC-COs separated clearly into two distinct clusters in a PCA plot (**Fig. 4c**). Differential gene expression analysis showed striking induction of chemokine gene expression, including for *IL1A, CXCL8*, *CXCL6, CXCL11*, and *IL1B*, yet with no detectable levels of *IFN-* I or *IFN-III*, which is consistent with recent reports^8–10^ (**Fig. 4d**). Ingenuity Pathway Analysis of the differential gene expression list highlighted the production of nitric oxide and reactive oxygen species, oxidative phosphorylation, as well as IL-15 production (**Fig. 4e**). The hPSC-COs were pre-treated with MPA or QNHC and infected with a relatively high titer of SARS-CoV-2 virus (MOI=0.1). Immunostaining confirmed the decrease of SARS-CoV-2^+^ cells in MPA or QNHC-treated hPSC-COs (**Fig. 4f**). Finally, western blotting assays confirmed the ability of MPA and QNHC to block SARS-CoV-2 infection of Vero cells (**Extended Data Fig. 4**).

**Figure 4.**
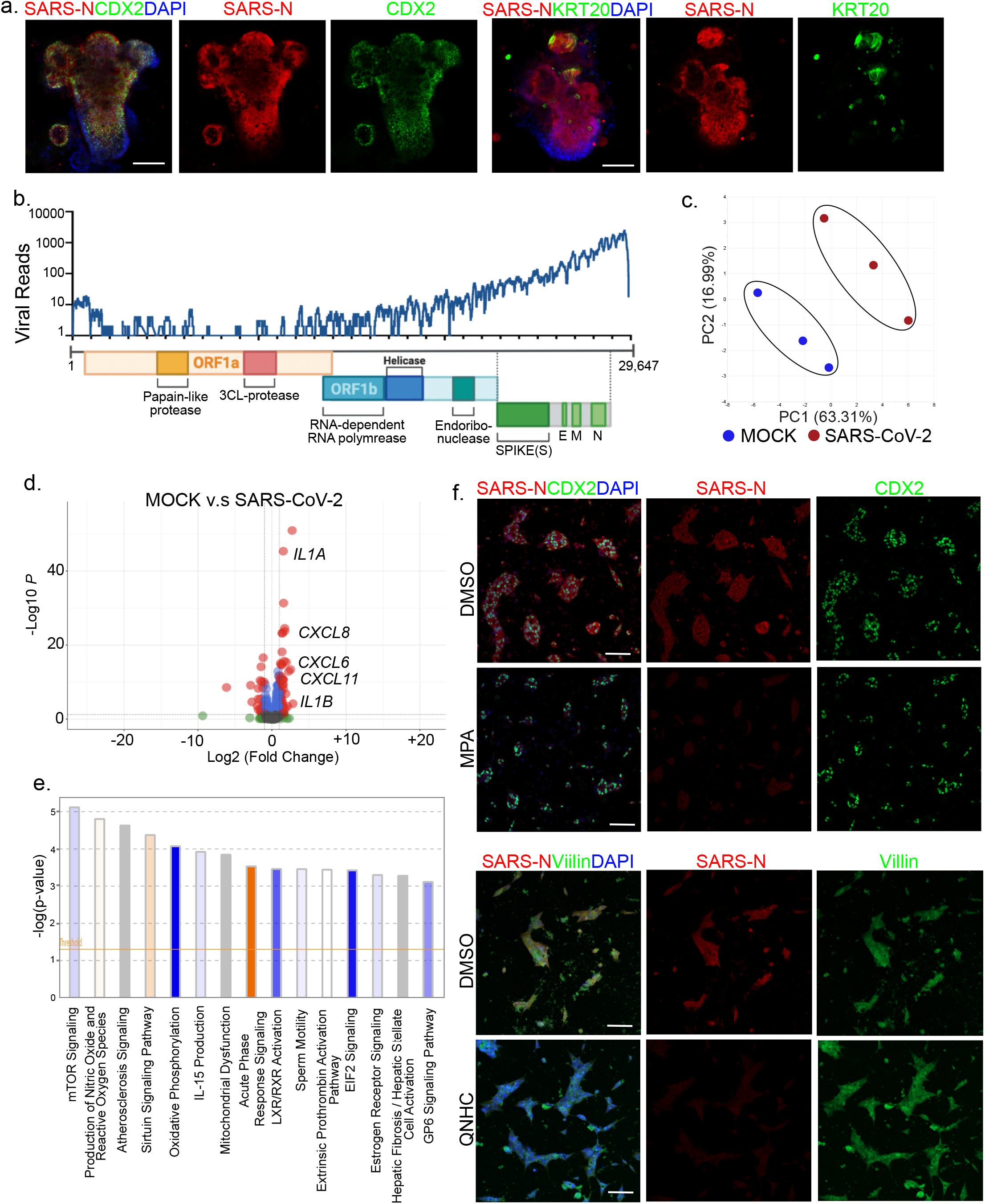
Quinacrine dihydrochloride and mycophenolic acid block the infection of SARS-CoV-2 virus. **a,** Immunofluorescent staining to detect SARS-CoV-2-nucleocapsid protein (SARS-N) in COs co-stained for CDX2 or KRT20. DAPI stains nuclei. Scale bar = 100 μm. **b,** Alignment of the transcriptome with the viral genome in SARS-CoV-2 infected hPSC-COs. Schematic noted of the SARS-CoV-2 genome. **c,** PCA plot of gene expression profiles from MOCK-infected and SARS-CoV-2 infected hPSC-COs at 24 hpi (MOI=0.1). **d,** Volcano plots indicating differentially expressed genes in hPSC-COs comparing MOCK and SARS-CoV-2 infected hPSC-COs at 24 hpi (MOI=0.1). Differentially expressed genes (p-adjusted value < 0.05) with a log_2_ (Fold Change) > 2 are indicated in red. Non-significant differentially expressed genes with a log_2_ (Fold Change) > 2 are indicated in green. **e,** IPA of differentially expressed genes in d. **f,** Immunofluorescent staining of SARS-CoV-2 nucleocapsid protein (SARS-N) of 3 μM MPA, 4.5 μM QNHC or DMSO-treated hPSC-COs at 24 hour post-SARS-CoV-2 infection (MOI=0.1). Scale bar = 100 μm.

In summary, we report that hPSC-derived COs express ACE2 and TMPRS2S2 and are permissive to SARS-CoV-2 infection. There is currently a lack of physiologically relevant models for COVID-19 disease that enable drug screens. Previous studies were based on clinical data or transgenic animals, for example mice that express human ACE2. However, such transgenic animals fail to fully recapitulate the cellular phenotype and host response of human cells^11,12^. We adapted a hPSC-derived CO platform for high throughput drug screening. Using disease-relevant normal colonic human cells, we screened 1280 FDA-approved compounds and identified MPA and QNHC, two drugs that can block the entry of SARS-CoV-2 into human cells. Strikingly, in this assay, the efficacies of MPA and QNHC for blocking viral entry are more than 5 times higher than chloroquine, a drug recently authorized by the FDA for emergency use to treat COVID-19 patients. Moreover, the MPA concentrations effective in blocking viral entry and replication are below that which is routinely used in clinical therapy^13^. MPA is a reversible, non-competitive inhibitor of inosine-5′-monophosphate dehydrogenase and is used widely and safely as an immunosuppressive drug (mycophenolate mofetil; CellCept) to prevent organ rejection after transplantation and for the treatment of autoimmune diseases^14^. MPA has been reported to block replication of human immunodeficiency virus^15^, dengue^16^, as well as Middle East respiratory syndrome coronavirus (MERS-CoV)^17^. Several studies on MERS-CoV suggest that MPA may noncompetitively inhibit the viral papain-like protease while also altering host interferon response^17,18^. A recent study also predicts MPA would modulate the interaction between host protein inosine-5’-monophosphate dehydrogenase 2 (IMPDH2) and SARS-CoV-2 protein nonstructural protein 14 (nsp14)^19^. Furthermore, a clinical study of MERS-CoV suggested that the patients treated with mycophenolate mofetil has 0% mortality rate, which is significantly lower than the overall mortality rate as 37%^20^. QNHC (Acriquine^®^, Atabrine^®^, Atebrin^®^, Mepacrine^®^) is an FDA-approved antimalarial drug used more recently as an anthelmintic, antiprotozoal, antineoplastic agent, and antirheumatic^21^. Recent studies have shown that quinacrine protects mice against Ebola virus infection *in vivo ^22^*. Both MPA and QNHC can be considered candidates for clinical trials of COVID-19 therapy.

**Extended Data Figure. 1.**
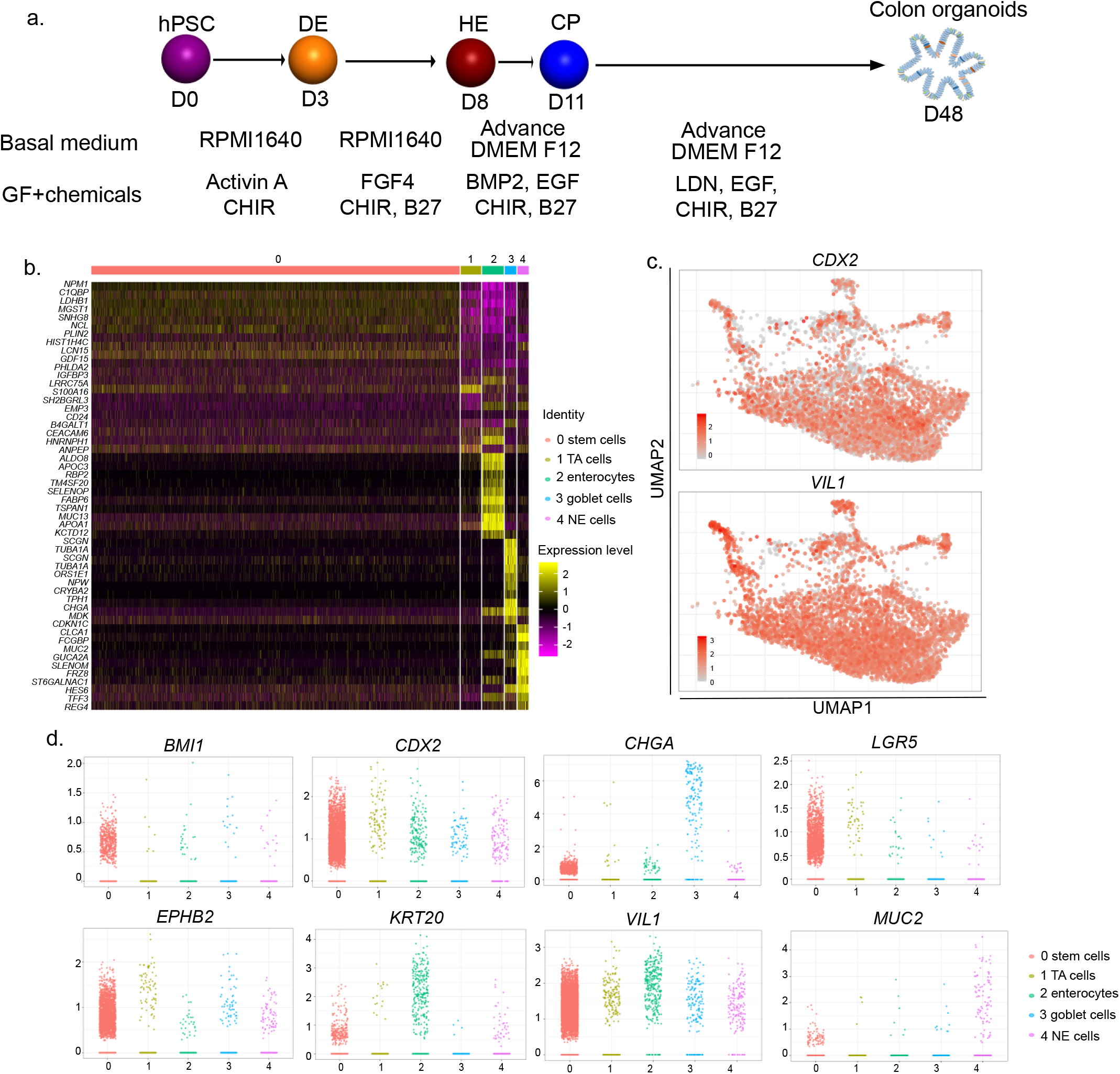
Single cell RNA-seq analysis of hPSC-derived COs. **a**, Schematic of protocol and conditions for hPSC differentiation to generate COs. **b**, Heatmap of top 10 differentially expressed genes in each cluster of single cell RNA-seq data. **c**, UMAP of *CDX2* and *VIL1*. **d**, Jitter plots for expression levels of colonic markers. Related to Fig. 1.

**Extended Data Figure. 2.**
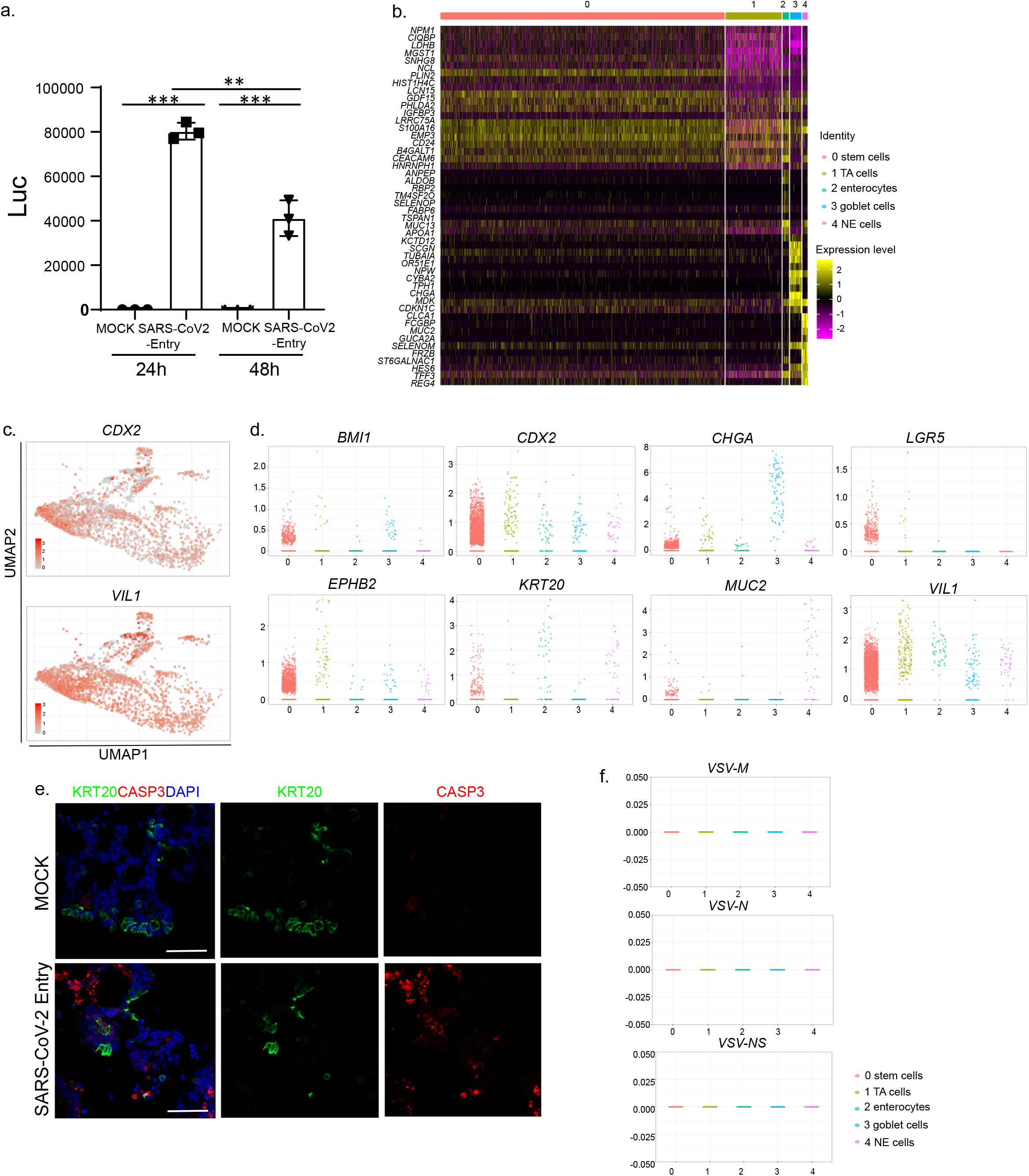
Single cell RNA-seq analysis of hESC-derived COs at 24 hours post infection with SARS-CoV-2 pseudo-entry virus. **a**, Relative luciferase levels in lysates derived from hPSC-derived COs inoculated with pseudo-entry virus at 24 or 48 hpi (MOI=0.01). **b**, Heatmap of top 10 differentially expressed genes in each cluster of single cell RNA-seq data. **c,** UMAP of *CDX2* and *VIL1*. **d**, Jitter plots for transcript levels of colonic markers. **e**, Representative immunostaining of infected COs co-stained for KRT20 and CASP3. Scale bar = 50 μm. **f**, Jitter plots of transcript levels for *VSV-M, VSV-N* and *VSV-NS* from hESC-derived COs without SARS-COV-2 infection (MOCK). Related to Fig. 2.

**Extended Data Figure. 3.**
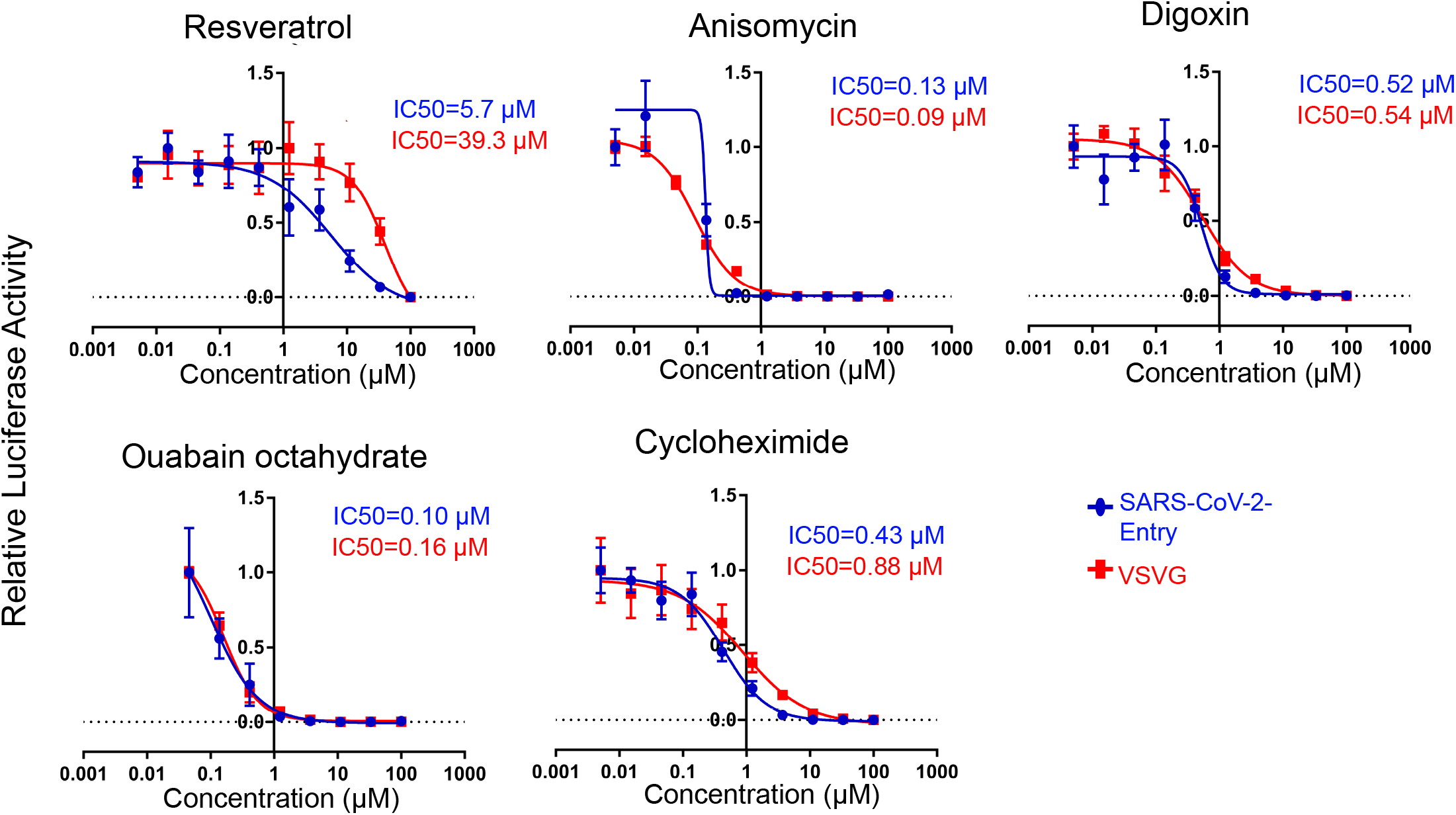
Inhibition curves of drug candidates on SARS-CoV-2 pseudo-entry virus compared to control VSVG virus infected hPSC-COs (For both viruses, MOI=0.01). Related to Fig. 3.

**Extended Data Figure. 4.**
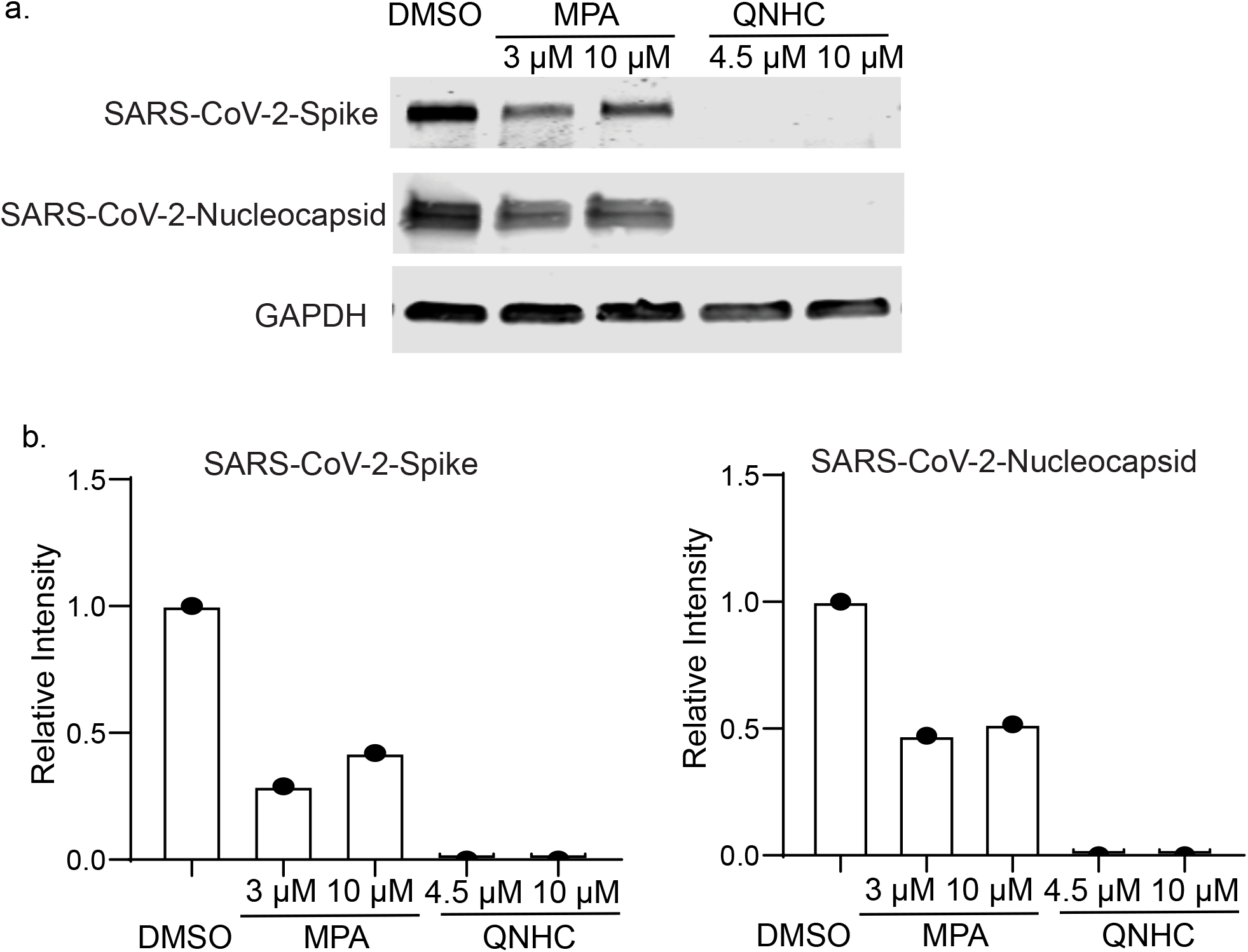
Western blotting (a) and quantification (b) to confirm MPA and QNHC’s anti-SARS-CoV-2 activity on Vero cells at 24 hpi (MOI=0.01). Related to Fig. 4.

## Methods

### hESC maintenance and colonic lineage differentiation

hESCs were grown and maintained on 1% Matrigel (Corning)-coated six-well plates in StemFlex medium (Gibco) at 37°C with 5% CO2. For definitive endoderm (DE) differentiation, hESCs were cultured to achieve 80-90% confluency, and treated with 3 μM CHIR99021 (CHIR, Stem-RD) and 100 ng/ml Activin A (R&D systems) in basal medium RPMI1640 (Cellgro) supplemented with 1X Pen-Strep (Gibco) for 1 day, and changed to the basal medium containing only 100 ng/ml Activin A the next day. To induce CDX2^+^ hindgut endoderm, DE were treated with 3 μM CHIR99021 and 500 ng/ml FGF4 (Peprotech) in RPMI1640 supplemented with 1X B27 supplement (Gibco) and 1X Pen-Strep (Gibco) for 4 days with daily changing of fresh media. Organoids began to bud out from the 2D culture during the hindgut differentiation process. The hindgut endoderm was then subjected to colonic lineage induction by treatment with 100 ng/ml BMP2 (Peprotech), 3 μM CHIR99021 and 100 ng/ml hEGF (Peprotech) in Advance DMEM F12 medium supplemented with 1X B27 supplement (Gibco), 1X GlutaMax (Gibco), 10 mM HEPES (Gibco) and 1X Pen-Strep (Gibco) for 3 days with daily changing of fresh medium. After colonic fate induction, the colon progenitor organoids were collected from the initial 2D cultures and embedded in a 100% Matrigel dome in a 24-well plate. Differentiation to mature colonic cell types was achieved by culturing these colon progenitor organoids in differentiation medium containing 600 nM LDN193189 (Axon), 3 μM CHIR99021 and 100 ng/ml hEGF in Advance DMEM F12 medium supplemented with 1X B27 supplement, 1X GlutaMax, 10 mM HEPES and 1X Pen-Strep. The differentiation medium was refreshed every 3 days for at least 40 days to achieve full colonic differentiation. The colon organoids were passaged and expanded every 10 – 14 days at 1:6 density. To passage the organoids, the Matrigel domes containing the organoids were scrapped off the plate and resuspended in cold splitting media (Advance DMEM F12 medium supplemented with 1X GlutaMax, 10 mM HEPES and 1X Pen-Strep). The organoids were mechanically dislodged from the Matrigel dome and fragmented by pipetting in cold splitting media. The old Matrigel and splitting media were removed after pelleting cells and the organoids were resuspended in 100% Matrigel. 50 μL Matrigel containing fragmentized colon organoids were plated in one well of a pre-warmed 24-well plate.

### Cell Lines

HEK293T (human [*Homo sapiens*] fetal kidney) and Vero E6 (African green monkey [*Chlorocebus aethiops*] kidney) were obtained from ATCC (https://www.atcc.org/). Cells were cultured in Dulbecco’s Modified Eagle Medium (DMEM) supplemented with 10% fetal bovine serum and 100 I.U./mL penicillin and 100 μg/mL streptomycin. All cell lines were incubated at 37°C with 5% CO_2_.

### SARS-CoV-2 Pseudo-Entry Viruses

Recombinant Indiana VSV (rVSV) expressing SARS-CoV-2 spikes were generated as previously described^23^. HEK293T cells were grown to 80% confluency before transfection with pCMV3-SARS-CoV2-spike (kindly provided by Dr. Peihui Wang, Shandong University, China) using FuGENE 6 (Promega). Cells were cultured overnight at 37°C with 5% CO2. The next day, medium was removed and VSV-G pseudotyped ΔG-luciferase (G*ΔG-luciferase, Kerafast) was used to infect the cells in DMEM at an MOI of 3 for 1 hr before washing the cells with 1X DPBS three times. DMEM supplemented with 2% fetal bovine serum and 100 I.U./mL penicillin and 100 μg/mL streptomycin was added to the infected cells and they were cultured overnight as described above. The next day, the supernatant was harvested and clarified by centrifugation at 300g for 10 min and aliquots stored at −80°C.

### SARS-CoV-2 Viruses

Severe acute respiratory syndrome coronavirus 2 (SARS-CoV-2), isolate USA-WA1/2020 (NR-52281) was deposited by the Center for Disease Control and Prevention and obtained through BEI Resources, NIAID, NIH. SARS-CoV-2 was propagated in Vero E6 cells in DMEM supplemented with 2% FBS, 4.5 g/L D-glucose, 4 mM L-glutamine, 10 mM Non-Essential Amino Acids, 1 mM Sodium Pyruvate and 10 mM HEPES. Infectious titers of SARS-CoV-2 were determined by plaque assay in Vero E6 cells in Minimum Essential Media supplemented with 2% FBS, 4 mM L-glutamine, 0.2% BSA, 10 mM HEPES and 0.12% NaHCO_3_ and 0.7% agar. All work involving live SARS-CoV-2 was performed in the CDC/USDA-approved BSL-3 facility of the Global Health and Emerging Pathogens Institute at the Icahn School of Medicine at Mount Sinai in accordance with institutional biosafety requirements.

### SARS-CoV-2 pseudo-entry virus infections

To assay pseudo-typed virus infection on colon organoids, COs were seeded in 24 well plates. Pseudo-typed virus was added at MOI=0.01 plus polybrene at a final concentration of 8 μg/mL, and the plate centrifuged for 1 hr at 1200g. At 3◻hpi, the infection medium was replaced with fresh medium. At 24◻hpi, colon organoids were harvested for luciferase assays or immunostaining analysis. For chemical screening analysis, colon organoids were digested by TrypLE and seeded in 384 well plates at 1×10^4^ cells per well. After chemical treatment, pseudo-typed virus was added at MOI=0.01 and the plate centrifuged for 1 hr at 1200g. At 24◻hpi, hPSC-COs were harvested for luciferase assays according to the Luciferase Assay System protocol (Promega).

### SARS-CoV-2 virus infections

hPSC-COs were infected with SARS-CoV-2 in the CO media at an MOI of 0.1, 0.05 or 0.01 as indicated and incubated at 37°C. At 24 hpi, cells were washed three times with PBS and harvested for either RNA analysis or immunofluorescence staining.

Approximately 2.5 × 10^5^ Vero E6 cells were pre-treated with the indicated compounds for 1 h prior to infection with SARS-CoV-2 at an MOI of 0.01 in DMEM supplemented with 2% FBS, 4.5 g/L D-glucose, 4 mM L-glutamine, 10 mM Non-Essential Amino Acids, 1 mM Sodium Pyruvate and 10 mM HEPES. At 24 hpi, cells were washed three times with PBS before harvesting for immunofluorescence staining or RNA or protein analysis. Cells were lysed in RIPA buffer for protein analysis or fixed in 5% formaldehyde for 24 h for immunofluorescent staining, prior to safe removal from the BSL-3 facility.

### Colon organoid processing and immunostaining

The colon organoids were released from Matrigel using Cell Recovery Solution (Corning) on ice for 1 hr, followed by fixation in 4% paraformaldehyde for 4 hr at 4°C, washed twice with 1X PBS and allowed to sediment in 30% sucrose overnight. The organoids were then embedded in OCT (TissueTek) and cryo-sectioned at 10 μm thickness. For indirect immunofluorescence staining, sections were rehydrated in 1X PBS for 5 min, permeabilized with 0.2% Triton in 1X PBS for 10 min, and blocked with blocking buffer containing 5% normal donkey serum in 1X PBS for 1 hr. The sections were then incubated with the corresponding primary antibodies diluted in blocking buffer at 4°C overnight. The following day, sections were washed three times with 1X PBS before incubating with fluorophore-conjugated secondary antibody for one hr at RT. The sections were washed three times with 1X PBS and mounted with Prolong Gold Antifade mounting media with DAPI (Life technologies). Images were acquired using an LSM880 Laser Scanning Confocal Microscope (Zeiss) and processed with Zen or Imaris (Bitplane) software.

### Immunofluorescent staining

Organoids and tissues were fixed in 4% PFA for 20 min at RT, blocked in Mg^2+^/Ca^2+^ free PBS plus 5% horse serum and 0.3% Triton-X for 1 hr at RT, and then incubated with primary antibody at 4°C overnight. The information for primary antibodies is provided in **Extended Data Table 2**. Secondary antibodies included donkey anti-mouse, goat, rabbit or chicken antibodies conjugated with Alexa-Fluor-488, Alexa-Fluor-594 or Alexa-Fluor-647 fluorophores (1:500, Life Technologies). Nuclei were counterstained by DAPI.

### Western blot

Protein was extracted from cells in Radioimmunoprecipitation assay (RIPA) lysis buffer containing 1X Complete Protease Inhibitor Cocktail (Roche) and 1X Phenylmethylsulfonyl fluoride (Sigma Aldrich) prior to safe removal from the BSL-3 facility. Samples were analysed by SDS-PAGE and transferred onto nitrocellulose membranes. Proteins were detected using rabbit polyclonal anti-GAPDH (Sigma Aldrich, G9545), mouse monoclonal anti-SARS-CoV-2 Nucleocapsid [1C7] and mouse monoclonal anti-SARS-CoV-2 Spike [2B3E5] protein (a kind gift by Dr. T. Moran, Center for Therapeutic Antibody Discovery at the Icahn School of Medicine at Mount Sinai). Primary antibodies were detected using Fluorophore-conjugated secondary goat anti-mouse (IRDye 680RD, 926-68070) and goat anti-rabbit (IRDye 800CW, 926-32211) antibodies. Antibody-mediated fluorescence was detected on a LI-COR Odyssey CLx imaging system and analyzed using Image Studio software (LI-COR).

### Single cell organoid preparation for scRNA-sequencing

The colon organoids cultured in Matrigel domes were dissociated into single cells using 0.25% Trypsin (Gibco) at 37°C for 10 min, and the trypsin was then neutralized with DMEM F12 supplemented with 10% FBS. The dissociated organoids were pelleted and resuspended with L15 Medium (Gibco) supplemented with 10 mM HEPES, and 10 ng/ml DNaseI (Sigma). The resuspended organoids were then placed through a 40 μm filter to obtain a single cell suspension, and stained with DAPI followed by sorting of live cells using an ARIA II flow cytometer (BD Biosciences). The live colonic single cell suspension was transferred to the Genomics Resources Core Facility at Weill Cornell Medicine to proceed with the Chromium Single Cell 3’ Reagent Kit v3 (10x Genomics, product code # 1000075) using 10X Genomics Chromium Controller. A total of 10,000 cells were loaded into each channel of the Single-Cell A Chip to target 8000 cells. Briefly, according to manufacturer’s instruction, the sorted cells were washed with 1x PBS + 0.04% BSA, counted by a Bio-Rad TC20 Cell Counter, and cell viability was assessed and visualized. A total of 10,000 cells and Master Mixes were loaded into each channel of the cartridge to generate the droplets on Chromium Controller. Beads-in-Emulsion (GEMs) were transferred and GEMs-RT was undertaken in droplets by PCR incubation. GEMs were then broken and pooled fractions recovered. After purification of the first-strand cDNA from the post GEM-RT reaction mixture, barcoded, full-length cDNA was amplified via PCR to generate sufficient mass for library construction. Enzymatic fragmentation and size selection were used to optimize the cDNA amplicon size. TruSeq Read 1 (read 1 primer sequence) was added to the molecules during GEM incubation. P5, P7, a sample index, and TruSeq Read 2 (read 2 primer sequence) were added via End Repair, A-tailing, Adaptor Ligation, and PCR.

The final libraries were assessed by Agilent Technology 2100 Bioanalyzer and sequenced on Illumina NovaSeq sequencer with pair-end 100 cycle kit (28+8+91).

### Sequencing and gene expression UMI counts matrix generation

T FASTQ files were imported to a 10x Cell Ranger - data analysis pipeline (v3.0.2) to align reads, generate feature-barcode matrices and perform clustering and gene expression analysis. In a first step, cellranger *mkfastq* demultiplexed samples and generated fastq files; and in the second step, cellranger count aligned fastq files to the reference genome and extracted gene expression UMI counts matrix. In order to measure viral gene expression, we built a custom reference genome by integrating the four virus genes and luciferase into the 10X pre-built human reference (GRCh38 v3.0.0) using cellranger *mkref*. The sequences of four viral genes (VSV-N VSV-NS, VSV-M and VSV-L) were retrieved from NCBI (https://www.ncbi.nlm.nih.gov/nuccore/335873), and the sequence of the luciferase was retrieved from HIV-Luc.

### Single-cell RNA-seq data analysis

We filtered cells with less than 300 or more than 8000 genes detected as well as cells with mitochondria gene content greater than 30%, and used the remaining cells (6175 cells for the uninfected sample and 2962 cells for the infected sample) for downstream analysis. We normalized the gene expression UMI counts for each sample separately using a deconvolution strategy^24^ implemented by the R scran package (v.1.14.1). In particular, we pre-clustered cells in each sample using the *quickCluster* function; we computed size factor per cell within each cluster and rescaled the size factors by normalization between clusters using the *computeSumFactors* function; and we normalized the UMI counts per cell by the size factors and took a logarithm transform using the *normalize* function. We further normalized the UMI counts across samples using the *multiBatchNorm* function in the R batchelor package (v1.2.1). We identified highly variable genes using the *FindVariableFeatures* function in the R Seurat (v3.1.0)^25^, and selected the top 3000 variable genes after excluding mitochondria genes, ribosomal genes and dissociation-related genes. The list of dissociation-related genes was originally built on mouse data^26^, we converted them to human ortholog genes using Ensembl BioMart. We aligned the two samples based on their mutual nearest neighbors (MNNs) using the *fastMNN* function in the R batchelor package, this was done by performing a principal component analysis (PCA) on the highly variable genes and then correcting the principal components (PCs) according to their MNNs. We selected the corrected top 50 PCs for downstream visualization and clustering analysis. We ran the uniform manifold approximation and projection (UMAP) dimensional reduction using the *RunUMAP* function in the R Seurat^25^ package with training epochs setting to 2000. We clustered cells into eight clusters by constructing a shared nearest neighbor graph and then grouping cells of similar transcriptome profiles using the *FindNeighbors* function and *FindClusters* function (resolution set to 0.2) in the R Seurat package. We identified marker genes for each cluster by performing differential expression analysis between cells inside and outside that cluster using the *FindMarkers* function in the R Seurat package. After reviewing the clusters, we merged four clusters that were likely from stem cell population into a single cluster (*LGR5^+^* or *BMI1^+^* stem cells) and kept the other four clusters (*KRT20^+^* epithelial cells, *MUC2^+^* goblet cells, *EPHB2^+^* TA cells, and *CHGA^+^* NE cells) for further analysis. We re-identified marker genes for the merged five clusters and selected the top 10 positive marker genes per cluster for heatmap plot using the *DoHeatmap* function in the R Seurat package^25^.

### In vivo transplantation and drug evaluation

hPSC-COs were harvested by cell scraper, mixed with 20 μl Matrigel (Corning) and transplanted under the kidney capsule of 7-9 weeks old male NSG mice. Two weeks post-transplantation, SARS-CoV-2 pseudo-entry virus was inoculated locally at 1×10^3^ FFU. At 24 hpi, the mice were euthanized and used for immunohistochemistry analysis.

To determine the MPA’s activity *in vivo*, the mice were treated with 50 mg/kg MPA in (10%DMSO/90% corn oil) by IP injection. Two hours after drug administration, SARS-CoV-2 pseudo-entry virus was inoculated locally at 1×10^3^ FFU. At 24 hpi, the mice were euthanized and used for immunohistochemistry analysis.

All animal work was conducted in agreement with NIH guidelines and approved by the WCM Institutional Animal Care and Use Committee (IACUC) and the Institutional Biosafety Committee (IBC).

### High throughput chemical screening

To perform the high throughput small molecule screen, hPSC-COs were dissociated using TrypLE for 20 min in a 37℃ waterbath and replated into 10% Matrigel-coated 384-well plates at 20,000 cells/40 μl medium/well. After 6 hr, cells were treated with compounds from an in-house library of ~1280 FDA-approved drugs (Prestwick) at 10 μM. DMSO treatment was used as a negative control. One hour late, cells will be infected with SARS-CoV-2 pseudo virus (MOI=0.01). After 24 hpi, hPSC-COs were harvested for luciferase assay following the Luciferase Assay System protocol (Promega).

### RNA-Seq following viral infections

Organoid infections were performed at an MOI of 0.1 and harvested at 24 hpi in DMEM supplemented with 0.3% BSA, 4.5 g/L D-glucose, 4 mM L-glutamine and 1 μg/ml TPCK-trypsin. Total RNA was extracted in TRIzol (Invitrogen) and DNase I treated using Direct-zol RNA Miniprep kit (Zymo Research) according to the manufacturer’s instructions. RNA-seq libraries of polyadenylated RNA were prepared using the TruSeq RNA Library Prep Kit v2 (Illumina) or TruSeq Stranded mRNA Library Prep Kit (Illumina) according to the manufacturer’s instructions. cDNA libraries were sequenced using an Illumina NextSeq 500 platform. Raw reads were aligned to the human genome (hg19) using the RNA-Seq Aligment App on Basespace (Illumina, CA), following differential expression analysis using DESeq2^27^. Differentially expressed genes (DEGs) were characterized for each sample (p adjusted-value < 0.05). Volcano plots were constructed using custom scripts in R.

## Statistical analysis

N=3 independent biological replicates were used for all experiments unless otherwise indicated. n.s. indicates non-significance. *P*-values were calculated by unpaired two-tailed Student’s t-test unless otherwise indicated. \**p*<0.05, \*\**p*<0.01 and \*\*\**p*<0.001.

## Acknowledgements

This work was supported by the Department of Surgery, Weill Cornell Medicine (T.E., F.P, S.C.), by the Defense Advanced Research Projects Agency (DARPA) through a contract with B.T. (DARPA-16-35-INTERCEPT-FP-006) and by the Jack Ma Foundation to D.D.H. The authors would like to thank Dr. Tom Moran, Center for Therapeutic Antibody Discovery at the Icahn School of Medicine at Mount Sinai for providing anti-SARS-Cov-SPIKE antibody. We are also very grateful for technical support and advice provided by Lee Cohen-Gould and Robert Lance Furler in the Cell Screening Core Facility of WCM.

## Author contributions

S. C., T. E., F. P., D. H., R.S, H. J. C., H. W., conceived and designed the experiments. X. D., Y.H., L. Y., F. P., performed organoid differentiation, *in vivo* transplantation, pseudo-virus infection and drug screening.

P. W, Y. Hu., performed SARS2-CoV-2 pseudo-entry virus related experiments.

B. N., and B. T., performed SARS2-CoV-2 related experiments.

T. Z., J. X. Z., D. X., X. W., performed the scRNA-sequencing and bioinformatics analyses.

## Competing interests

The authors declare the following competing interests: R.E.S. is on the scientific advisory board of Miromatrix Inc. The other authors have no competing of interest.

## Data Availability

scRNA-seq and RNA-seq data is available from the GEO repository database, accession number GSE147975.

---

**Extended Data Table. 1.**
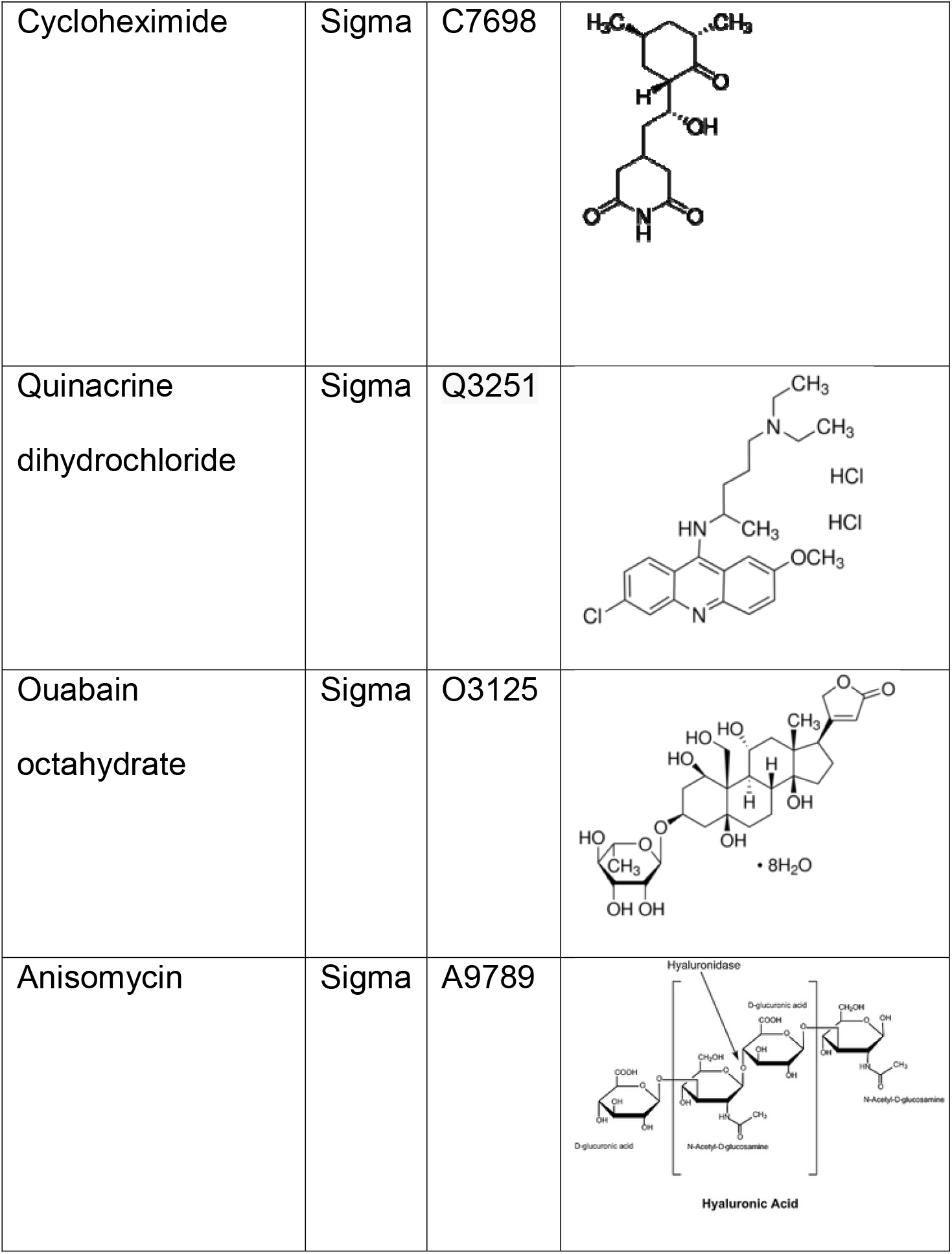
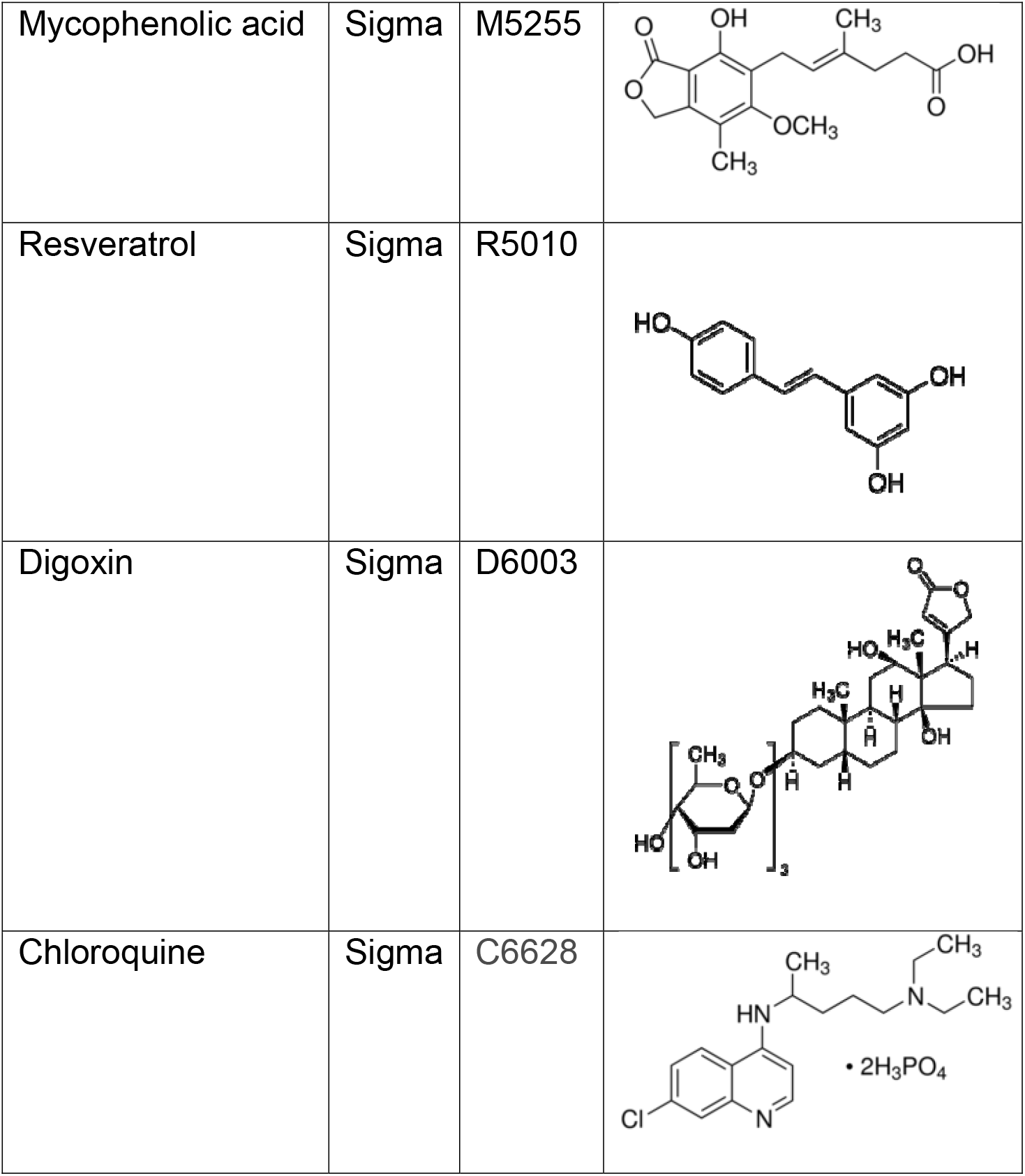
Primary hit compounds from chemical screening.

|  |  |  |
| --- | --- | --- |
| Cycloheximide | Sigma | C7698 |
| Quinacrine<br>dihydrochloride | Sigma | Q3251 |
| Ouabain<br>octahydrate | Sigma | O3125 |
| Anisomycin | Sigma | A9789 |
| Mycophenolic acid | Sigma | M5255 |
| Resveratrol | Sigma | R5010 |
| Digoxin | Sigma | D6003 |
| Chloroquine | Sigma | C6628 |

**Extended Data Table. 2.** Antibodies used for immunostaining.

|  | Species | Vendor | Catalogue number | dilution |
| --- | --- | --- | --- | --- |
| **Primary antibodies** |  |  |  |  |
| CDX2 | Mouse | Biogenex | MU392A-UC | 1:500 |
| Villin | Goat | Santa Cruz | sc-7672 (C-19) | 1:200 |
| Villin | Mouse | Santa Cruz | sc-58897 | 1:200 |
| Chromogranin A | Rabbit | Immunostar | 20086 | 1:300 |
| SATB2 | Mouse | Sigma | HPA001042 | 1:50 |
| Cytokeratin-20 | Mouse | Santa Cruz | sc-56522<br>(SPM140) | 1:100 |
| Mucin2 | Rabbit | Abcam | ab76774 | 1:100 |
| EPHB2 | Goat | R&D | AF467 | 1:100 |
| ACE2 | Goat | R&D | AF933 | 1:500 |
| ACE2 | Rabbit | Abcam | ab15348 | 1:500 |
| Firefly luciferase | Mouse | Thermo Fisher | 35-6700 | 1:200 |
| Firefly luciferase | Rabbit | Abcam | Ab185924 | 1:100 |
| Anti-SARS-CoV<br>Nucleocapsid | Rabbit | Rockland | 200-402-A50 | 1:250 |
| Cleaved Caspase-3<br>(Asp175) | Rabbit | Cell Signaling | 9661S | 1:200 |
| **Secondary antibodies** |  |  |  |  |
| Cy3-conjugated anti<br>rabbit | Donkey | Jackson<br>Immunoresearch | 711-165-152 | 1:500 |
| Cy3-conjugated anti<br>goat | Donkey | Jackson<br>Immunoresearch | 705-165-003 | 1:500 |
| Cy3-conjugated anti<br>mouse | Donkey | Jackson<br>Immunoresearch | 715-165-150 | 1:500 |
| Cy2-conjugated anti<br>rabbit | Donkey | Jackson<br>Immunoresearch | 711-225-152 | 1:500 |
| Cy2-conjugated anti<br>goat | Donkey | Jackson<br>Immunoresearch | 705-225-147 | 1:500 |
| Cy2-conjugated anti<br>mouse | Donkey | Jackson<br>Immunoresearch | 715-225-150 | 1:500 |
| Donkey anti-Mouse<br>IgG Secondary<br>Antibody, Alexa Fluor<br>488 | Donkey | Thermo Fisher | A21202 | 1:1000 |
| Donkey anti-Rabbit<br>IgG Secondary<br>Antibody, Alexa Fluor<br>555 | Donkey | Thermo Fisher | A31572 | 1:1000 |
| Donkey anti-Goat IgG Secondary Antibody, Alexa Fluor 555 | Donkey | Thermo Fisher | A32816 | 1:1000 |
| Donkey anti-Rabbit IgG Secondary Antibody, Alexa Fluor 647 | Donkey | Thermo Fisher | A32795 | 1:1000 |

